# Persistence of genetically engineered canola populations in the U.S. and the adventitious presence of transgenes in the environment

**DOI:** 10.1101/2023.12.05.570248

**Authors:** Steven Travers, D. Bryan Bishop, Cynthia Sagers

**Author notes:** Corresponding author (CS). These authors contributed equally to the work (Steven Travers, Cynthia Sagers).

## Abstract

Feralization of genetically engineered (GE) crops increases the risk that transgenes will become integrated into natural and naturalizing plant populations. A key assumption of the management of GE crops is that populations of escaped plants are short-lived and that the risks they pose, therefore, are limited. However, few populations of escaped crop plants have been tracked over the long term so our understanding of their persistence in ruderal or natural landscapes is limited. In 2021, we repeated a large-scale road survey of feral GE canola populations in North Dakota, USA, initially conducted in 2010. Our objectives were to determine the current distribution of feral canola populations, and to establish the relative frequency of GE and non-GE phenotypes in populations of canola throughout North Dakota, where most of the canola in the USA is grown. Our results indicate that, although the incidence of feral canola was less in 2021 than 2010, escaped canola populations remain common throughout the state. The prevalence of alternate forms of GE herbicide resistance changed between surveys, and we found an overabundance of non-GE plants compared to the frequency of non-transgenic forms in cultivation. Indirect evidence that canola escaped from cultivation has established persistent populations outside of management includes finding plants with multiple transgenic traits, and locating populations far from transportation routes. We conclude that feral canola populations expressing transgenic herbicide resistance are established outside of cultivation, that they may be under selection for loss of the transgene, but that they nonetheless pose long-term risks by harboring transgenes in the unmanaged landscape.

## Introduction

The adventitious presence of transgenes in production systems continues to pose significant risks to U.S. agriculture. Detection of transgenes in foodstuffs has disrupted international and domestic markets, stalled supply chains, and created mistrust among consumers [1]. Durisin and Wilson [2] estimated that the threat of contamination of conventional and organic crops by genetically engineered (GE) varieties cost producers $6.3B US when food companies and foreign markets rejected transgene-contaminated supplies. Limiting the movement of transgenes in agricultural systems is central to reducing market losses, but such efforts are frustrated by the biology of plants and their ability to out-maneuver containment strategies. Numerous crop species have escaped cultivation and established persistent populations outside of cultivation [3]. Outside of managed croplands, escaped crops may evolve rapidly to become more closely adapted to, and tolerant of the non-agronomic environment [4]. Once established, they constitute a gene pool capable of generating weedy pests that carry the additional risks of harboring beneficial transgenes [5]. The risks of an adventitious presence, then, are enhanced when GE crop species escape cultivation to form stable, naturalized populations in the unmanaged landscape.

De-domestication, or feralization, of GE crops increases the risk that transgenes will escape cultivation and integrate with wild populations of plants, including close relatives. De-domestication is an evolutionary process by which domesticated plants or animals escape intensive management by humans to form independent reproducing populations [3]. De-domesticates are known to originate from within crop species through multiple routes: through mutation and selection (endo-ferals), by crossing between distinct populations or land races (exo-endo ferals), or by hybridizing with wild relatives (exoferals) [3]. Once established, the genetic architecture of de-domesticated populations may be further shaped by hybridization and introgression among wild, feral, and domesticated forms [6, 7, 8]. Wild traits, such as seed-shattering, asynchronous flowering and seed dormancy, are at times quickly recovered by de-domesticates as a product of selection in non-agronomic habitats [4]. The resulting feral populations, now adapted to local environments and tolerant of competition, are rich targets for ongoing gene flow from agricultural fields [9]. When they compete with related crops, or introduce deleterious traits to commercial fields, de-domesticates pose a threat to the integrity and profitability of commercial production systems.

Among domesticated species, kohl crops have an evolutionary history punctuated with repeated bouts of feralization, hybridization and introgression [8, 10, 11]. *Brassica napus* (2n = 4x = 38, AACC), an oilseed crop, is an allotetraploid originating from natural hybridization and genome duplication events between *B. rapa* (turnip) and *B. oleracea* (cabbage) [12, 13, 14, 15]. A detailed domestication history of *B. napus* remains somewhat unresolved, however, since all “wild” populations of *B. napus* appear to be feralized [15, 16, 17]. It is believed that *Brassica napus* originated 7,180 – 1,910 YBP from the Mediterranean basin or some agricultural regions such as northern or western Europe where *B. rapa* and *B. oleraceae* coexisted [16, 18]. Since the origins of *B. napus*, repeated introgressions have occurred both from *B. rapa* to *B. napus*, and *B. napus* to *B. rapa* [19, 20]. Further, recent population structure analysis of *B. napus* verifies ongoing hybridization with *B. rapa* and *B. oleracea* and the persistence of conspecific alleles over the last 1000 years [15, 17, 21]. Ongoing gene flow is evidenced by contemporary commercial varieties of *B. napus* that spontaneously hybridize with *B. rapa* and other *Brassica* species in crop fields and non-agronomic habitats [9, 22].

Canola (Brassica napus L.)(2n = 4n = AACC) is an industrial hybrid of *B. rapa* (2n = 20, AA) and *B. oleracea* (2n = 18, CC). It has become one of the most important oilseed crops worldwide and has been cultivated extensively for more than 100 years [23]. It is thought to have been domesticated relatively recently (within the last 300–400 years) [16]. Populations of domesticated canola growing outside of cultivation have been reported from Belgium, Austria, Denmark, France, Germany, U.K., Australia, the Netherlands, and New Zealand [1]. “Wild” traits still expressed in commercial canola, such as seed shattering, may contribute to its ready escape from cultivation [1]. Moreover, canola seeds retain partial dormancy and may remain viable in the soil seedbank for up to three years [24, 25]. The combined effects of seed loss on harvest and seed dormancy rapidly stock the soil seed bank, contributing to the high incidence of volunteer canola in and around agricultural fields [26, 27].

The U.S. was an early adopter of genetically engineered (GE) canola in the mid-1990s and now nearly all U.S. canola is GE [28]. In 2011, we described the presence of escaped GE herbicide resistance (GE HR) canola populations North Dakota, USA, where most of the U.S. canola is grown. More than 75% of the plants sampled from roadside populations were GE HR and most populations contained a mixture of GE HR phenotypes, either glyphosate resistance or glufosinate resistance [29]. The source of these populations has yet to be determined, but it was generally believed they result from seed spill during transportation [30]. We repeated the survey in 2021 with the objectives to:

- Determine the current distribution of feral canola populations in North Dakota, USA
- Establish the relative frequency of GE HR traits in North Dakota populations
- Investigate the distribution and frequency of non-GE plants in roadside samples

Between census periods, U.S. production of canola increased substantially, with a reported increase in the frequency of glufosinate cultivation. Therefore, if seed spill is the dominant factor maintaining feral populations, as has been argued [30], we anticipate an increase in the number and size of feral populations, and a change in the phenotypic profile of feral populations.

## Methods

In 2021, we conducted systematic roadside surveys to quantify the presence and abundance of feral GE and non-GE canola populations in North Dakota, USA, beginning 14 June 2021 and continuing through 7 July 2021. No permits or approvals were required for this work as all samples were collected in the public right of way and no protected species were sampled. We adopted methods similar to the survey of 2010 [29]. Field crews established transects on major east-west highways throughout the state. A 2×50 m quadrat was established every 8.05 km (5 miles) of roadway on one or both sides of the road, where traffic permitted, in which all identifiable *B. napus* plants were counted. We drove a total of 6373 km and sampled 62.5 km of roadside habitat (1.0% of the distance driven). Sampling was conducted early in the summer prior to the onset of flowering of cultivated canola. When canola was present at a sampling site, one randomly selected plant was collected, photographed and archived as a voucher specimen. Leaf fragments from voucher specimens were tested for the presence of CP4 EPSPS protein (confers tolerance to glyphosate herbicide) and PAT protein (confers tolerance to glufosinate herbicide) with TraitChek™ immunological lateral flow test strips (Romer Labs, Inc., Newark, DE). Previous studies have demonstrated the utility of the lateral flow strips in detecting the expression of transgenes from field samples [29. 31, 32]. Test strips are not available for a third, non-GE resistance trait, resistance to Clearfield™ herbicide, which is reported to comprise less than 5% of the canola grown in the region (R Beneda, pers comm). At random intervals, single plants were tested with multiple test strips to assure that test results were repeatable and reliable. No failures were detected during the course of the study. To determine if populations of escaped canola are composed of multiple genotypes, multiple plants were sampled at nine randomly selected, large feral canola populations and tested for the presence of CP4 EPSPS or PAT proteins. Finally, to confirm a reported shift from CP4 EPSPS+ to PAT+ cultivation, we collected canola growing immediately outside of 58 agricultural fields and presumably established by seed spill, and tested for the expression of GE HR..

## Results

### 2021

Populations of escaped canola were once again widespread throughout the state (Fig 1). In 2021 canola was present at 42% (262/623) of the road survey sampling sites. Of those, 76% (199/262) expressed at least one transgene: 67% (176/262) were positive for only PAT (glufosinate resistance); 8% (20/262) positive for only CP4 EPSPS (glyphosate resistance); and 1.1% (3/262) expressed both forms of herbicide resistance. Densities of canola plants at collection sites ranged from 0 to 10 plants m^−2^ with an average of 0.4 plants m^−2^.

**Fig 1.**
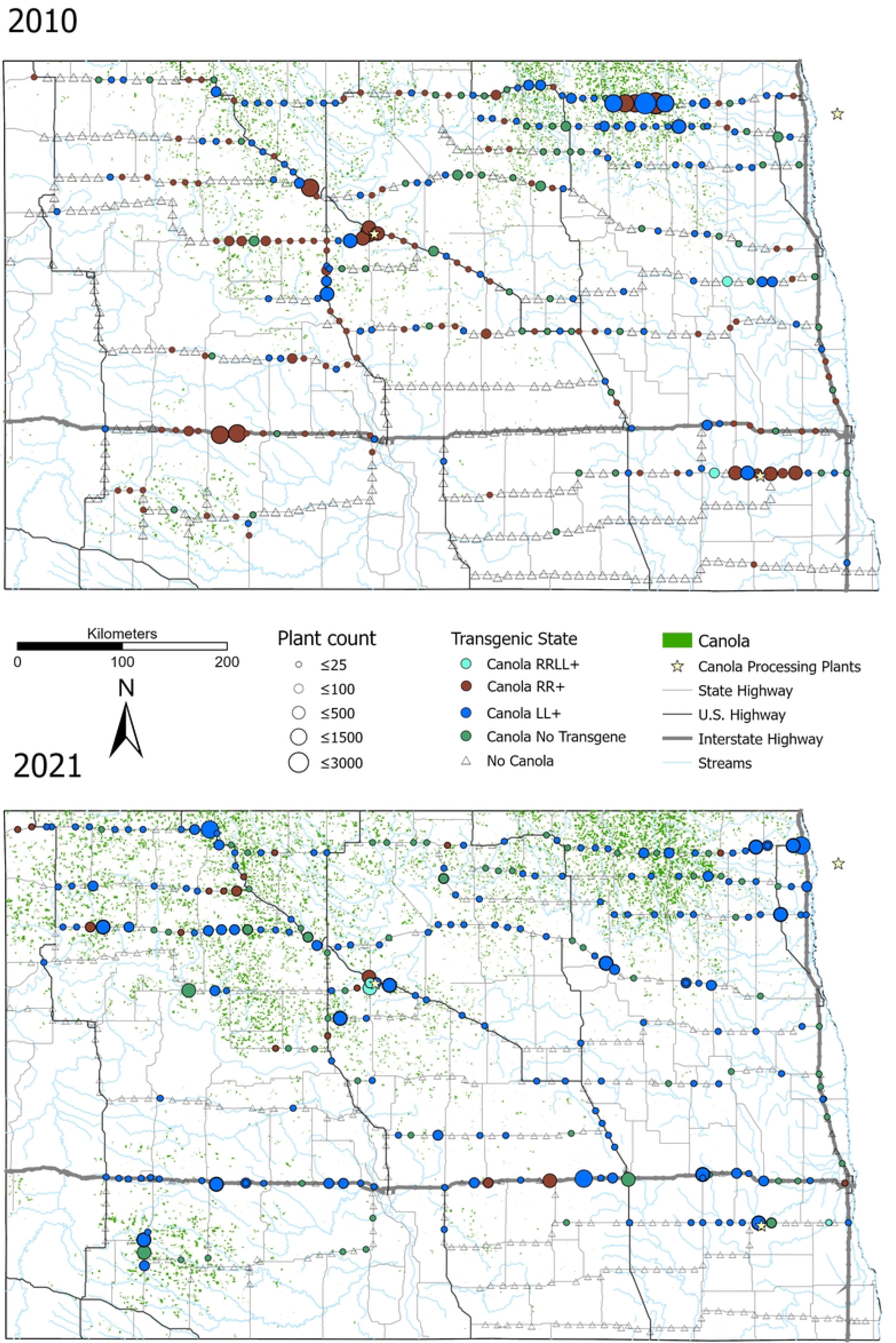
Results of 2010 and 2021 field surveys of feral canola populations in North Dakota. Solid black lines indicate major highways. Collection sites indicated by symbols: triangles = no canola; circles = canola present. Shading indicates transgenic character state: blue = PAT+; red = CP4 EPSPS+; turquoise = PAT+/CP4 EPSPS+; green = non-GE. Circle diameter reflects estimate of local population size. Green stippling indicates locations of canola production fields in 2009 (panel A) and 2020 (panel B). Oilseed processing plants are noted by stars. Survey data from Schafer et al. 2011 [29] and Travers et al. (this study). Sources for GIS data layers are available in Appendix I.

Sampling within large populations of feral canola revealed a mix of both herbicide resistant -, and non-GE phenotypes. In 2022, we sampled 25 plants from large, escaped populations and found that all populations in this sample consisted of a mix of resistance phenotypes (CP4 EPSPS+, PAT +, CP4 EPSPS+/PAT+, non-GE).

In 2021 we sampled canola from 58 fields and found that all samples expressed GE herbicide resistance: 90% PAT+; 9% CP4 EPSPS+; 2% containing both transgenes. We found evidence of neither non-GE nor Clearfield™ canola cultivation in our sample.

### 2021 vs. 2010

In 2021 we sampled a total of 623 roadside sites, 1% fewer than in 2010 (Fig 2). Overall, we encountered far fewer plants in the survey in 2021 than in 2010 (9884 vs. 18,960, respectively), despite an increase of 26% in the total number of acres of cultivated canola [33]. The proportion of sampling sites with canola dropped somewhat in 2021 to 42% (262/623) from 45% (286/631). We detected a significant shift in the relative frequency of HR phenotypes with PAT becoming the dominant form of HR (67% in 2021 versus 49% in 2010); CP4 EPSPS becoming less frequent (10% in 2021 versus 51% in 2010), and non-GE increasing slightly to 24% (63/262) from 20% (57/286) (Fig 2). Overall, the changes between census periods in the frequency of GE HR and non-GE phenotypes in feral populations was significant (χ^2^ =81.5, df = 2, p <.001).

**Fig 2.**
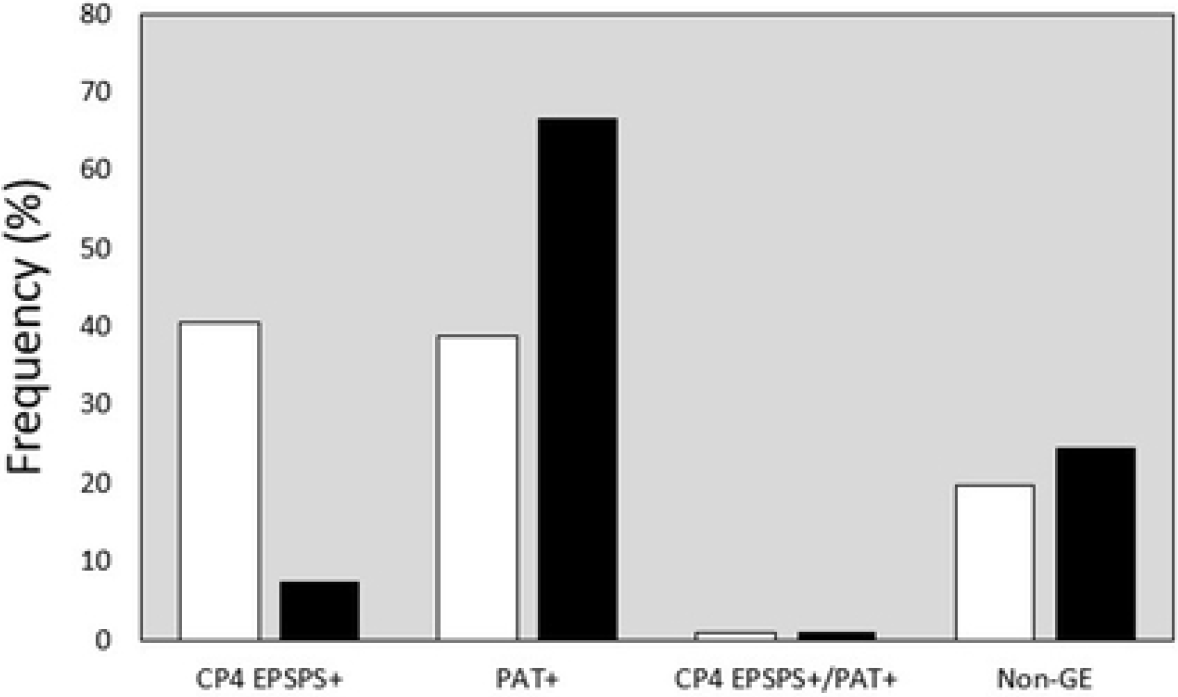
Frequency of GE and non-GE samples in 2010 and 2021 field surveys of feral canola in North Dakota. Open bars: 2010 survey; closed bars: 2021 survey. CP4 EPSPS+ - glyphosate resistant; PAT+ - glufosinate resistant. Changes in relative frequencies are significant (χ^2^ =81.5, df = 2, p <.001).

In both census periods, populations of transgenic canola were most dense along major transport routes, at construction sites and in regions of intense canola cultivation (Fig. 1). At a finer scale, feral populations appeared denser at junctions between major roadways, access points to crop fields and bridges, and at intersections of roadways with railway crossings. At these sites, seed spill during transport appears the most likely mechanism for the escape of transgenic canola. Nonetheless, feral canola plants found occasionally at remote locations far from canola production, transportation, or processing facilities in 2010 were encountered more frequently in 2021. In 2021, in addition to growing in road verges (Fig 3A) and construction sites (Fig 3B), we discovered populations in riparian habitats (Fig 3C) and natural areas, including two populations in protected areas (Sheyenne National Grassland, Upper Souris National Wildlife Refuge). Moreover, canola plants growing at streamside locations were robust and phenotypically more similar to crop canola than feral plants in road verges and construction fill (Fig 3C).

**Fig 3.**
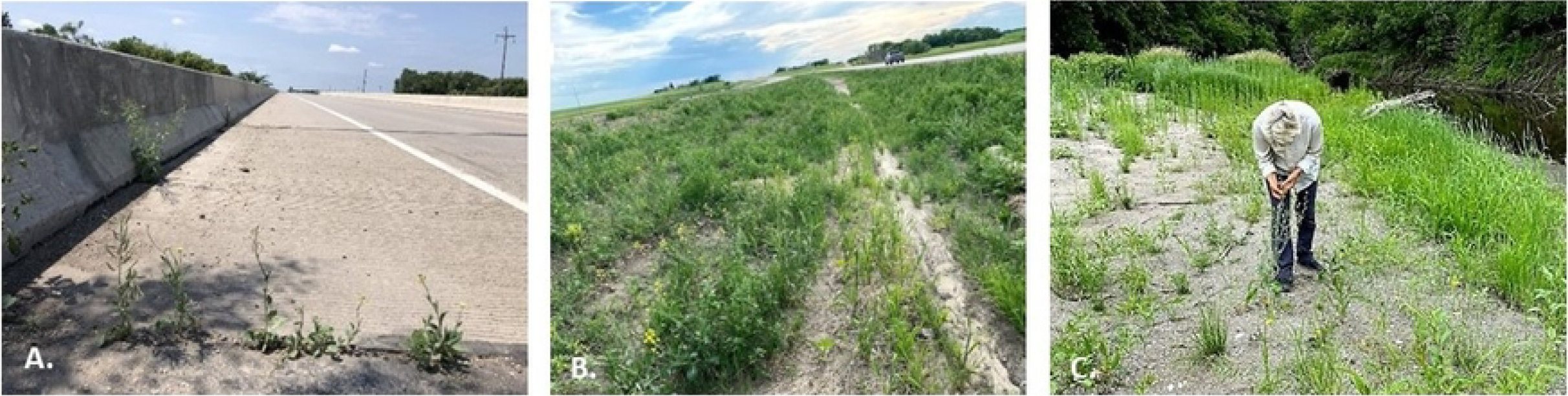
Feral canola in non-agronomic environments: (A) Along transportation routes (State Hwy 46, Richland Co., North Dakota, 2021); (B) In soil fill-dirt at road construction sites (Hwy I-94, 41 mi west of Jamestown, Kidder County, North Dakota, 2021), (C) In riparian habitats (Sheyenne River, near the Sheyenne National Grassland, Ransom Co, North Dakota, 2021). GE canola was detected at each site.

## Discussion

### Long term trends in feral canola populations in North Dakota

Feral canola remains widespread along the roadsides of North Dakota. The total number of feral canola plants in the sample, and the frequency of populations fell between 2010 and 2021 sampling periods. The incidence of GE HR populations growing outside of cultivation decreased, despite a large increase in the number of cultivated acres (+26%). A drop in occurrence despite an increase in cultivated acres may be explained by severe drought conditions that persisted through the growing season of 2021: May through August 2021 rainfall recorded in the centrally located city of Minot, ND, in 2021 was 38% below average [34].

The make-up of feral populations changed substantially over the course of a decade. Whereas feral plants in 2010 were equally likely to be resistant to glyphosate (CP4 EPSPS+) as glufosinate (PAT+), by 2021 glufosinate resistance was nine times more common. We would expect to see this rate of change if the following conditions hold: a) feral populations are ephemeral, b) escaped populations are maintained through seed spill, and c) cultivation had shifted strongly to PAT+ from CP4 EPSPS. Although USDA no longer publishes data on GE HR canola acreages, a limited survey of ag fields in 2021 showed a predominance of PAT+ varieties (52/58 fields). Further, conversation with grower’s associations revealed PAT+ to be more popular with growers than CP4 EPSPS+ in recent years (Barry Coleman, pers. comm.). This may have resulted from an increasing number of glyphosate-resistant weeds that evolved in the intervening decade. Currently in the U.S., 17 weed species are resistant to glyphosate whereas only three species are resistant to glufosinate [35]. In addition, large populations of feral canola express a mixture of resistance phenotypes, consistent with repeated immigration events. Together, these findings support the idea that escaped populations are dynamic and persist under selection and with repeated re-introduction of new seeds each growing season. Our use of sampling sites along transportation routes undoubtedly created bias toward detecting the effects of seed spill, however. Future studies of population dynamics, structure, and origins should sample further afield, including riparian areas and grasslands.

### Long-term persistence of feral canola

Movement by transport may explain the current distribution of feral GE canola along transportation routes in North Dakota, but it does not explain the incidence of feral non-GE plants. From 2010 to 2021, the frequency of non-GE canola escaped from cultivation increased to 24.0% from 19.9%, despite the near universal cultivation of GE HR canola in North Dakota. The most recent published estimate of GE HR canola is 95% of cultivated acres [28]. In our small sample of canola fields in 2021, none were non-GE, however. The high (20-24%) frequency and persistence of non-GE forms in our sample could be due to several factors:1) non-transgenic canola (e.g., Clearfield™, Westar varieties) may have a selective advantage in non-agronomic habitats, 2) transgenes carry a cost and are lost by natural selection in non-agronomic environments, 3) non-GE plants could be the products of older feral lineages, escaped cultivation before the release of GE canola, or, 4) non-GE canola is from much older feral lineages now widely distributed in North America [36]. Regardless of its origins, the high incidence of non-GE forms argues for the longer term persistence of feral *B. napus* in the environment and suggests that the dynamics of feral GE canola are more complex than initially believed. Future studies of the genetic structure of escaped populations could easily resolve questions of origins, but progress to this end has been slowed by the difficulties of genomic analysis of the tetraploid genome of *B. napus*. Nevertheless, a handful of studies have made headway untangling the genomics and structure of feral *B. napus* populations [37, 38, 39].

## Conclusions

These findings raise compelling questions about the dynamics of feral GE canola populations. A decadal change in transgenic genotype frequencies in feral populations may simply reflect a change in the types of GE canola crops planted over this time [41]. Data on GE planted acreages are not immediately available to test this idea. If so, it would solidify the argument that populations in our sample are largely maintained by seed spill. The high frequency of non-GE forms in the unmanaged landscape (20-25%) relative to non-GE planted acreages (0-5%) is far more compelling. Their sources are unknown, but these populations represent a unique opportunity to understand the histories of populations in the process of de-domestication and the mechanisms of transgene persistence in them. They may be recent introductions, or they may represent long-term populations that have persisted in the landscape before the introduction of commercial canola. As such, they merit further analyses at the genomic and population level. Fully understanding the risks posed by the adventitious presence of transgenes will require further genomics studies of populations growing in naturalized habitats away from transportation routes in areas, such as North Dakota, where GE canola production is widespread [40].

There is now evidence for self-sustaining feral canola populations in 14 countries on five continents [reviewed in 1]). In Europe and Japan, plants in feral populations are similar genetically to varieties planted in fields nearby but nonetheless show a degree of divergence [37, 38, 39, 41]. Feral populations have consistently been found to have higher levels of genetic diversity and unique alleles and combinations of alleles, suggesting the admixture of wild, cultivated and feral genomes, multiple incidences of escape, and long-term persistence of feral populations [37, 39]. Although genetically engineered canola is reported growing outside of cultivation in six countries: USA [29], Japan ([39, 42], Canada [43, 44], Switzerland [45], Argentina [46], and the Netherlands [47], the detailed population analyses required to assess their risks are few and restricted to regions where commercial GE canola production is limited.

## Appendix I. GIS data sources

